# Constraining inference of across-region interactions using neural activity perturbations

**DOI:** 10.1101/2025.11.20.689584

**Authors:** Diya Basrai, JeongYoon Lee, Amy Kristl, Peiyu Wang, Andrew Miri, Joshua I. Glaser

## Abstract

Functional interactions between brain regions are often inferred from multi-region models fit to neural activity recorded in behaving animals. Here, we show that inference of across-region interactions is hindered by the wide breadth of model fits consistent with naturally occurring neural activity. In contrast, models fit to activity that includes region-wide activity perturbations provide well-constrained estimates of across-region interactions; simulations suggest these estimates can be accurate.

## Main

Understanding neural system function requires characterizing functional interactions within networks of neurons across the brain. Models are commonly trained to infer these interactions by learning connection strengths that best reconstruct naturally occurring neural activity patterns recorded in behaving animals^1–3^. While this approach is increasingly applied to quantify interactions between brain regions from multi-region recordings^4–6^, the space of well-fitting model parameters may be quite large^7–9^, which would render across-region interactions ambiguous. Instead of fitting models solely to naturally occurring neural activity, theoretical results suggest models can be better constrained by fitting to activity patterns that include the effects of activity perturbations^10–12^. We therefore examined how well multi-region recordings alone constrain estimates of across-region interactions, and whether including region-wide perturbations will better constrain these estimates.

We first sought to compare estimates of across-region influence from models fit to naturally occurring activity alone with those from fits to activity affected by rapid optogenetic perturbations. We simultaneously recorded neural activity in two connected regions and sporadically silenced activity in one. We used Neuropixels 1.0 arrays to record activity in mouse forelimb M1 and M2 while vGAT-ChR2-eYFP mice performed a self-paced naturalistic climbing task^13,14^. Here mice perform a broad range of limb movements to traverse an unpredictable terrain, avoiding the movement stereotypy that can build with experience in other motor behavioral paradigms^15,16^. During climbing bouts, we briefly and sporadically applied 25-ms pulses of blue light to activate inhibitory interneurons in either region; equivalent behavioral events were notated to serve as controls (Fig. 1a,b). In the directly inactivated (upstream) region, light pulses rapidly suppress activity in the putative pyramidal neurons that mediate across-region interactions^13,17^, leading to subsequent suppression of activity in the other (downstream) region (Fig. 1c).

**Figure 1.**
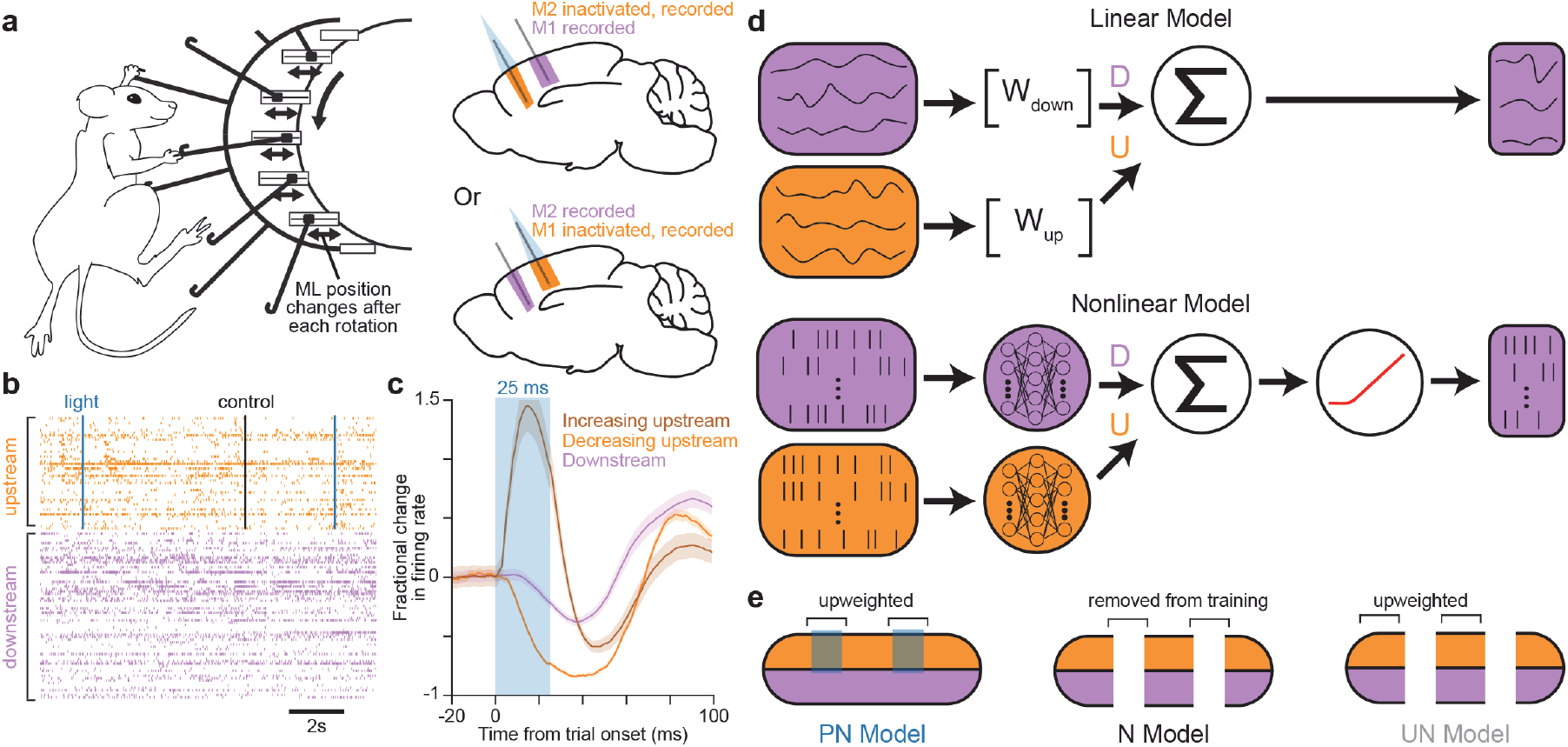
Paradigm to assess model fitting with and without perturbation. **a**, Schematic depicting climbing behavior (left) and mouse brain schematics depicting simultaneous M1 and M2 recordings while inactivating one region (right). M2 was inactivated in 6 sessions from 2 mice and M1 was inactivated in 7 sessions from 2 different mice. **b**, Example spike raster with the time of perturbation and control trials indicated. Control trials, where no light was applied, were used to define held-out segments of naturally occurring activity used for model testing. **c**, Mean ± SEM fractional change in firing rates averaged across perturbation trials from all 13 sessions. Increasing and decreasing upstream neurons were distinguished by the effects of perturbation on their trial averaged firing rates. **d**, Schematic of linear (top) and nonlinear (bottom) model architectures. **e**, Schematic of activity time series used as training data for each model type, including upweighted and removed segments.

We next trained models to predict activity in the downstream region during active climbing from the preceding activity in both the upstream and downstream regions. Importantly, our models made predictions by combining the output of distinct downstream and upstream networks, allowing separation of within-region (downstream; *D*) and across-region (upstream; *U*) contributions (Fig. 1d). We employed a linear model that predicted the top 20 principal components (PCs) for activity in the downstream region from linear combinations of downstream and upstream PCs. We also employed a nonlinear model that predicted single-neuron firing rates in the downstream region by combining outputs from downstream and upstream feedforward neural networks (FNNs), which received the firing rates in their respective regions as inputs.

We then could directly compare across-region contributions from models fit to naturally occurring activity alone (N models) and models fit to both naturally occurring activity and perturbation responses (PN models). Because perturbation responses constituted a small fraction of recorded activity, we heavily weighted these responses when measuring PN model performance (Fig. 1e). To control for this upweighting, we also fit models on naturally occurring activity alone with additional upweighting of some randomly selected activity segments (UN models). Models fit only to perturbation responses predicted naturally occurring activity relatively poorly and were not considered further.

To determine how well naturally occurring activity and perturbation responses constrain estimates of across-region interactions, we quantified the range of across-region contributions that yielded model performance close to the best fit. We fit models while applying a range of regularization penalties separately to the weights of upstream and downstream activity components. This produced models spanning a continuum from negligible to strong across-region input. For each model, we quantified the relative contribution of regions (region quotient, RQ) as 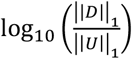. N models spanning a relatively wide range of RQs fit nearly equally well, indicating that naturally occurring activity is consistent with a broad range of upstream contributions (Fig. 2a). However, for PN models, there was a clear performance peak as RQ varied, indicating an unambiguous degree of upstream influence (Fig. 2b).

**Figure 2.**
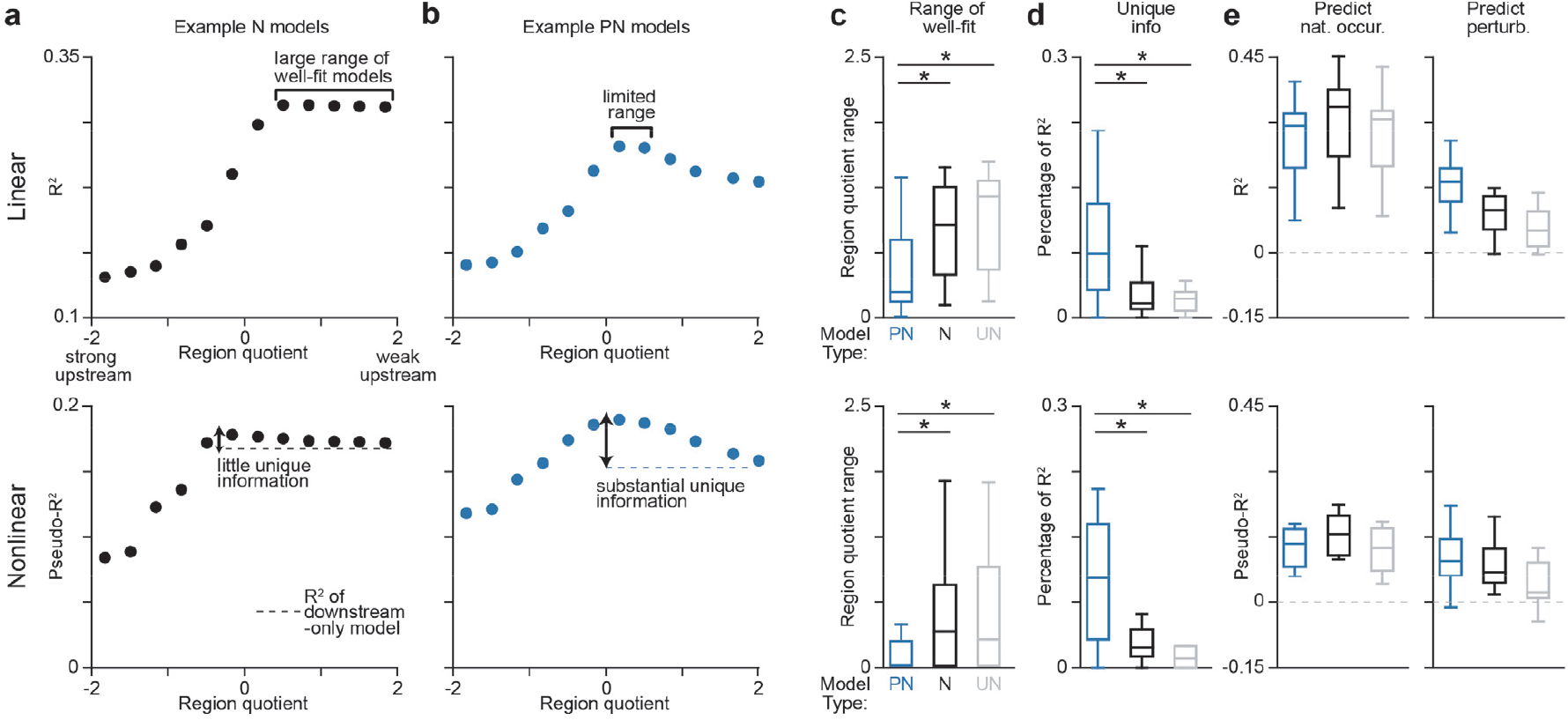
Inclusion of neural activity perturbations constrains estimates of across-region interactions. Top row: results for linear models. Bottom row: results for nonlinear models. **a**,**b**, Model performance versus region quotient for **(a)** N models and **(b)** PN models for one example session. Different region quotients are achieved via different regularization penalties on upstream and downstream weights. Model performance was measured only on the specific trial types the model was trained on; N and UN models were tested on held-out naturally occurring activity, and PN models were tested on both held-out naturally occurring and perturbation activity. For clarity, these plots show only a subset of all models fit. **c**, Box plots for the range of region quotient values for models within 1% of best-fit R^2^. For the linear models (top), significant differences were found between PN and N (p = 0.029, one-sided Wilcoxon rank-sum) and PN and UN (p = 0.029). For nonlinear models (bottom), significant differences were found between PN and N (p = 0.007), and PN and UN (p = 0.007). All box plots in this figure indicate the minimum, 1st quartile, median, 3rd quartile and maximum. Results in **c**,**d** use 324 (linear) or 30 (nonlinear) models fit with different regularization penalties. **d**, Box plots for the unique information provided by the upstream region, calculated by taking the fractional difference between best-fit model R^2^ and downstream-only model R^2^. For the linear models (top), significant differences were found between PN and N (p = 0.010, one-sided Wilcoxon rank-sum) and PN and UN (p = 0.010). For nonlinear models (bottom), significant differences were found between PN and N (p = 0.008), and PN and UN (p = 0.006). **e**, Model performance separated for held-out naturally occurring activity segments (left) and perturbations segments (right).

The range of models within 1% of peak performance (‘well-fit models’) was substantially and significantly lower for PN models compared to N and UN models (Fig. 2c). The broad range of well-fit N models suggests that in naturally occurring activity, the predictive information is concurrently present in both regions. To quantify the unique information present in the upstream region, we measured the fractional difference in predictivity between the best-fit model and a model that only uses downstream components. We found that the best-fit PN model relied on substantial unique information from upstream components. In contrast, N and UN models contained little unique information, such that models fit only to downstream activity performed nearly equally as well as models fit to activity in both regions (Fig. 2d). These results suggest that only models fit using perturbation responses can leverage unique information in the upstream region to identify a precise degree of across-region influence.

We then examined whether N and PN models differed in terms of their predictivity. We fit individual N and PN models using equal regularization penalties for both regions, as in standard model-fitting approaches. We compared the predictivity of the resulting models on the same held-out segments of naturally occurring activity and perturbation responses. PN models were able to better predict perturbation responses, while only slightly decreasing predictivity on naturally occurring activity. Thus the additional constraints did not greatly harm predictivity on naturally occurring activity (Fig. 2e). Furthermore, the greater predictivity of PN models on perturbation responses suggests that the best-fit PN models are meaningfully distinct from the best-fit N models. Thus perturbation responses have non-trivial effects on model structure.

While these results show that perturbation responses constrain models around a narrow best-fit functional connectivity between regions, its similarity to the actual underlying connectivity remains unclear. To address this, we examined connectivity inferred from simulations generated from a network model with known connectivity. We first fit multi-region recurrent neural networks (RNNs) to naturally occurring activity recorded in M1 and M2. We included the biological constraint that across-region weights in RNNs were positive and sparse (Fig. 3a). We then used the resulting models with known connectivity to run simulations. To mimic region-wide inactivation, we identified interneuron-like units in each region that did not make across-region connections and whose local connections were negatively weighted. During simulations, we sporadically drove these “interneurons” in the upstream region, causing short latency suppression of upstream projection neurons followed by activity suppression in the downstream region (Fig. 3b).

**Figure 3.**
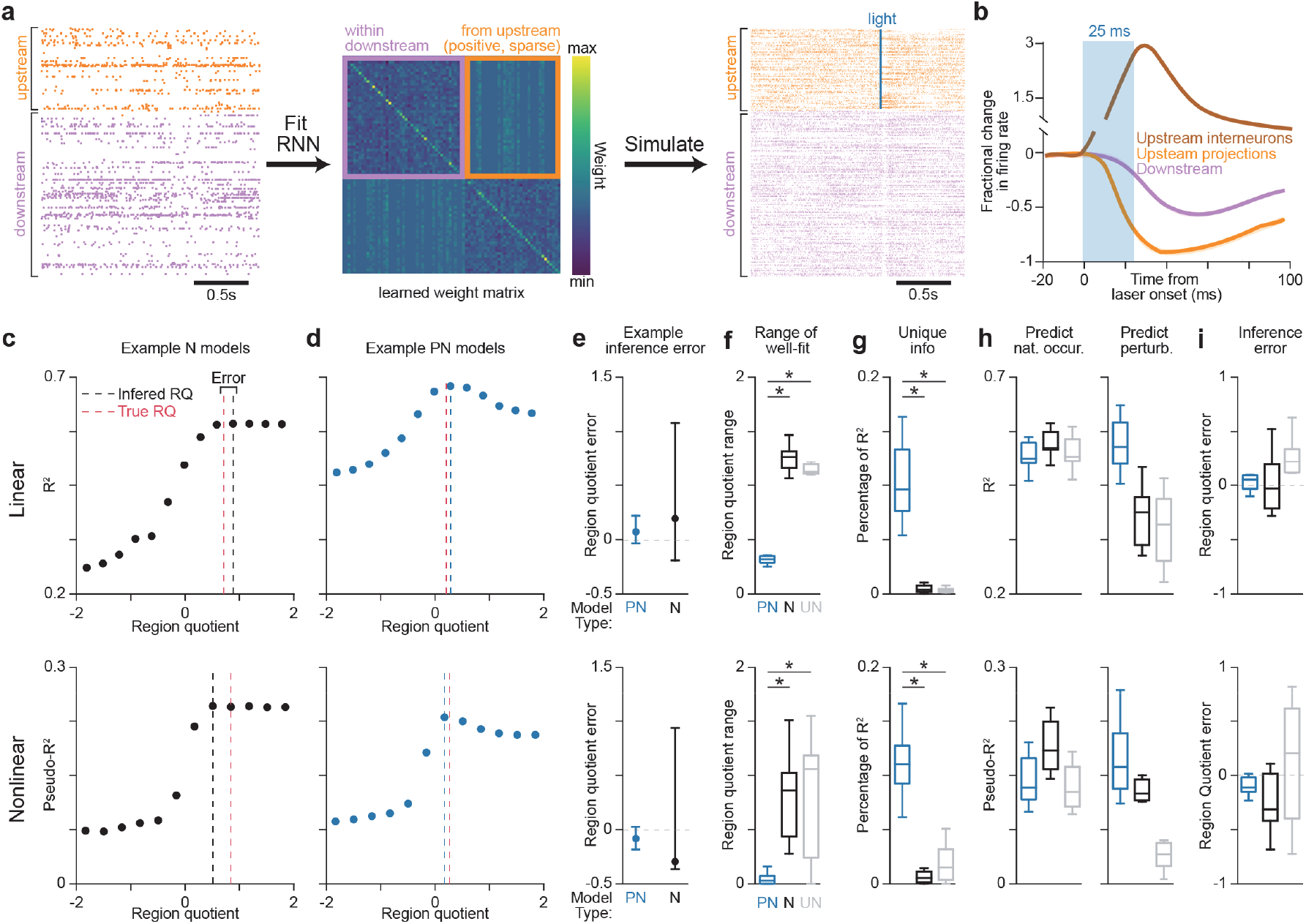
In simulations, perturbation responses constrain models around actual across-region influence. **a**, Schematic of simulation-based analysis. RNNs were fit for six sessions from three mice. **b**, Mean ± SEM fractional change in firing rates during simulated perturbation trials, averaged across all six RNNs. Upstream interneurons and upstream projections were defined by whether they had exclusively zero (interneurons) or nonzero (projections) weights on their connections to downstream neurons. **c-i**, Top row: results for linear models. Bottom row: results for nonlinear models. **c**,**d**, Model performance versus region quotient for N models (**c**) and PN models (**d**). Model performance was measured on held-out data. These plots show only a subset of all models fit for visualization purposes. All box plots in this figure indicate the minimum, 1st quartile, median, 3rd quartile and maximum. **e**, For one example RNN, whiskers depict the range of inference error for all well-fit models and filled circles depict the error of the best-fit model. Inference error is calculated from the difference between the inferred region quotient and the true region quotient. **f**, Box plots for the range of region quotient values for well-fit models. For the linear models (top), significant differences were found between PN and N ranges (p = 0.016, one-sided Wilcoxon rank-sum) and PN and UN ranges (p = 0.016). For nonlinear models (bottom), significant differences were found between PN and N ranges (p = 0.016), and PN and UN ranges (p = 0.039). Results in **f**,**g** use 384 (linear) or 30 (nonlinear) models fit with different regularization penalties. **g**, Box plots for the unique information provided by the upstream region, calculated by taking the fractional difference between best-fit model R^2^ and downstream-only model R^2^. For the linear models (top), significant differences were found between PN and N ranges (p = 0.016) and PN and UN ranges (p = 0.016). For nonlinear models (bottom), significant differences were found between PN and N ranges (p = 0.016), and PN and UN ranges (p = 0.016). **h**, Model performance separated for held-out naturally occurring activity segments (left), and perturbations trials (right). **i**, Box plots for the best-fit model inference error.

We first verified that the inclusion of perturbation responses in RNN simulations was necessary to find a constrained solution for across-region influence. As with actual recordings, we found well-fit N models with a broad range of RQs. In contrast, a single PN model emerged as the unambiguous best-fit model (Fig. 3c-e) with substantially more unique upstream information than N and UN models (Fig. 3f,g). Furthermore, when examining models fit using a standard regularization approach, PN models predicted perturbation responses significantly better than N models with only a slight reduction in predictivity on naturally occurring activity (Fig. 3h). Thus, even in a synthetic system where all neurons are sampled, perturbation responses meaningfully constrain estimates of across-region influence.

Finally, we asked whether including perturbation responses constrained models around the ground-truth connectivity. We calculated the inference error by taking the difference between the RQ of the best-fit model and the true RQ. The best-fit PN models had remarkably low inference error, tightly distributed around 0. With the true RQs ranging from 0.3 to 1.2, the interquartile range of inference errors was 0.12 for linear models and 0.13 for nonlinear models (Fig. 3e,i). In contrast, N and UN models had higher inference error; for N models, the interquartile range was 0.43 for linear models and 0.44 for nonlinear models. This demonstrates that including region-wide perturbations improves inference of across-region connectivity from simulated activity, yielding a close match to the ground truth. This suggests that models constrained by perturbation responses can effectively measure across-region interactions.

Our results demonstrate that the inclusion of responses to region-wide activity perturbations during model fitting enables precise estimates of underlying across-region interactions that are not achieved using naturally occurring activity alone. This approach would be complementary to, and could be combined with, other approaches that have recently shown promise for improving model accuracy, like including biological constraints (e.g., Dale’s Law) in model architecture^18,19^. Past observations that activity measurements leave functional connectivity ambiguous have attributed this deficit to the vast under-sampling of neurons and the limited range of observed activity states^20,21^. However, when fitting activity from RNN simulations where all neurons in the network were observed, we found that across-region interaction strengths estimated from unperturbed activity alone still remained ambiguous. Additionally, here we used activity recorded throughout a naturalistic climbing behavior that involves a broad range of limb movements. Despite the resulting exploration of a broader range of neural activity states than with traditional motor tasks^17,22^, perturbations were still necessary to well-constrain models.

## MATERIALS AND METHODS

### Experimental Animals

All experiments and procedures involving animals were performed in accordance with NIH guidelines and approved by the Institutional Animal Care and Use Committee of Northwestern University. A total of 14 adult male mice: 12 VGAT-ChR2-EYFP line 8 mice (B6.Cg-Tg(Slc32a1-COP4*H134R/EYFP) 8Gfng/J; Jackson Laboratories stock #014548) and 2 C57BL/6J mice (Jackson Laboratories stock #000664) were used in reported experiments and early pilot studies to establish methodology. All mice were individually housed under a 12 hr light/dark cycle during the course of the experiment. At the time of reported measurements, animals were 12-24 weeks old and weighed approximately 24-30 g. All animals were used for the first time in this experiment and were not previously exposed to pharmacological substances or altered diets.

### Climbing Apparatus

We used the climbing apparatus described in Koh et al^14^. Briefly, the climbing apparatus consisted of a 3D-printed cylindrical wheel with left handholds in fixed positions and right handholds connected to linear actuators that enabled their movement to random mediolateral positions after each time they rotate 180° past the mouse. This ensures that the sequence of right handhold positions is unpredictable. Mice are head-fixed in an upright position and pull on handholds with all four limbs to turn the wheel. The climbing is self-paced and not timed by any extrinsic sensory cues. Wheel position signals are recorded by an angular encoder, and water rewards are dispensed to mice by a lick tube positioned under the mouse’s mouth.

### Headplate Implantation and Behavior Training

Mice were outfitted with 3D printed plastic headplates and trained to climb as previously described^23^. Briefly, mice were anesthetized and a headplate was affixed to the skull using dental cement (Metabond, Parkell). After recovery from the headplate implantation, mice were placed on a restricted water schedule and acclimated to handling and head-fixation. After acclimation, mice were head-fixed on the wheel and encouraged to associate climbing with water rewards by the experimenter manually triggering water rewards during climbing movements. Once a mouse performed sustained climbing bouts (1-3 sessions, most typically 1), mice were trained using an automated script that dispensed water rewards after climbing bouts ended, with reward size adapted to incentivize increasingly long climbing bouts. After mice consistently engaged in climbing bouts (2-3 sessions), mice were prepared for recording sessions.

### Neural Recording and Inactivation

One day prior to the first recording session, mice were anesthetized and dental cement was removed from the skull to prepare either left forelimb M1 (fM1, centered 0.25 mm rostral and 1.5 mm lateral to bregma) or left forelimb M2 (fM2, centered 2.25 mm rostral and 1.00 mm lateral to bregma) for inactivation. The skull was thinned in a 2 mm diameter circle around the center of the site until translucent.

To expose both fM1 and fM2 for neural recording, two 0.5 mm craniotomies were drilled over both sites. A Pt-Ir reference wire soldered to a gold pin was implanted 1.5 mm deep in left posterior parietal cortex. A silicone elastomer (Kwik-Cast, World Precision Instruments) was applied to seal the recording craniotomies and surrounding skull, and the animal was allowed to recover overnight.

Prior to each recording session, the mouse was head-fixed on the wheel, the silicon elastomer was removed, and saline was applied to cover the craniotomies. Two Neuropixels 1.0 probes (Imec), pre-sharpened at a 20° angle, were inserted into fM1 and fM2 at a rate of 300 μm/min using 3-axis micromanipulators (M3-LS-3.4-15-XYZ, New Scale Technologies) to a final depth of 1.5 mm. Reference wires soldered onto each Neuropixel were connected to the reference pin. A 400 μm core, 0.50 NA optical patch cable terminating in a 2.5 mm ceramic ferrule (M128L01, Thorlabs) was positioned using an additional micromanipulator at a pre-calibrated distance to illuminate a spot of light (2 mm in diameter) over the thinned skull above fM1 or fM2. We used a 473 nm laser (MDL-III-473-200mW, Opto Engine LLC) to apply 25 ms pulses of blue light at an intensity of 10 mw/mm^2^. After Neuropixels were inserted and the patch cable’s ferrule was positioned, liquid paraffin was applied over the craniotomies and thinned skull to keep the brain moist over the ∼60-minute recording session.

During recording, a perturbation or control (no light) trial was occasionally triggered during active climbing provided the distance of wheel rotation during the current bout exceeded a random threshold between 0° and 20°. Trials were triggered no less than 3 s apart. A random 50% of trials were chosen to be controls. Experimental control signals were sampled at 4 kHz using a RHD2000 USB interface board and RHD USB interface software (Intan technologies). Neural recordings acquired at 30 kHz and bandpass filtered at 0.3 to 10 kHz using SpikeGLX software. At the end of each recording session, Neuropixels were retracted at a rate of 300 μm/min and silicon elastomer was applied to reseal the craniotomies. We performed 3-4 recording sessions with each mouse.

### Spike Sorting and Unit Curation

Prior to spike-sorting, raw activity along the length of the Neuropixels probe was first visualized on the Kilosort 3^24^ graphical interface. All channels outside the surface of the brain, clearly identifiable from a visually apparent change in voltage amplitude, were excluded. Spikes were then detected and sorted using Kilosort 3 with default parameters.

After sorting, we aligned each unit’s spike times to laser onset and excluded a small fraction of units in the illuminated region that had been assigned spikes that were likely light-induced electrical artifacts. These spikes were identifiable by spike times precisely at laser onset or laser offset on every trial, and were clearly distinguishable from light-activated interneurons due to absence of any sub-millisecond variability in spike timing across trials around laser onset.

### Data for Modeling

All models were fit to neural activity during climbing from individual recording sessions. From each session, we extracted 3 non-overlapping types of time series segments. Naturally occurring segments were defined using time windows during active climbing beginning at least 300 ms after light onsets (156-638 segments per session, median=236, segment length 99-6594 ms across sessions, median=1188 ms). Perturbation segments were defined as the -50 to 70 ms window around light onset times (58-244 segments per session, median = 82). For use as held-out naturally occurring activity for performance testing, we defined control segments from the -50 to 70 ms window around control trial onset times (55-275 segments per session, median = 81). We used segments including control trials as held-out naturally occurring activity to approximately match with perturbation segments, as perturbation and control trials were triggered based on the same criteria. Still, we obtained comparable results when using a more typical random sampling method for selecting time series to hold out from naturally occurring segments for performance testing.

In total, the ratio of the total number of timesteps in naturally occurring and perturbation segments was approximately 50:1. Given this imbalance, we heavily weighted timesteps from perturbation segments when measuring the performance of PN models. To account for this sample weighting, we additionally labeled randomly selected 120 ms windows within naturally occurring segments as ‘upweight segments,’ equal number to perturbation segments, which were upweighted in a separate set of models (UN models).

For fitting N and UN model parameters, training sets consisted of all naturally occurring segments. The PN training set consisted of all naturally occurring segments and a randomly sampled 80% of perturbation segments. For comparing model predictions on naturally occurring activity and perturbation responses (Fig. 2e and Fig. 3h), the naturally occurring test set consisted of all held-out control segments, and the perturbation activity test set consisted of a held-out 20% of perturbation segments. For analysis of model performance across different RQs, (Fig. 2a-d, and Fig. 3c-g), N and UN model test sets consisted of all held-out control segments. For PN models, the test set consisted of the 20% of held-out perturbation segments along with a randomly sampled 20% of control segments, reflecting the approximately 50/50 split of naturally occurring activity and upweighted perturbation activity in the training set.

### Models Overview

Linear models were fit to the principal components (PCs) for the firing rate time series of recorded neurons. Nonlinear models were fit to firing rate time series of individual neurons. We used both approaches in order to show robustness of our overall results. Below we refer to both the firing rates and PCs as ‘activity.’

Here, we briefly describe the general construction of inputs and outputs for both types of models. We constructed inputs ***X***_*d*_, ***X***_*u*_ and output ***Y*** that enabled autoregressive predictions of downstream activity (***Y***) from past activity in both the downstream (***X***_*d*_) and upstream (***X***_*u*_) region. Activity was averaged in 10 ms bins. For each timestep, we constructed input matrix ***X***_*d*_ and ***X***_*u*_ by concatenating the last 5 bins of activity from the downstream and upstream regions, respectively. We constructed the corresponding output matrix ***Y*** from the downstream activity at the subsequent timestep.

To illustrate ***X***_*u*_, ***X***_*d*_ and ***Y***, let ***M*** be a data matrix consisting of 1 downstream neuron and 1 upstream neuron with *t* timesteps:

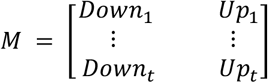

Then, ***X***_***d***_ and ***X***_***u***_, constructed from the past τ lags, will be:

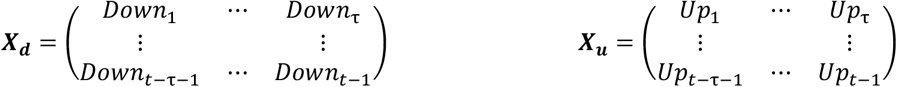

And the corresponding output ***Y*** will be:

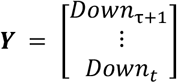

Both linear and nonlinear models predict ***Y*** by combining predictions from separate downstream (*D*) and upstream (*U*) networks, where *D* = *f*_*d*_(***X***_***d***_), and *U* = *f*_*u*_(***X***_***u***_).

### Linear Modelling

#### Model details

In our linear models, we predicted the top 20 downstream PCs from the previous activity of the top 20 downstream and upstream PCs. To construct the training set for each model, we first averaged activity in 10 ms bins. Then we applied PCA separately to each region, keeping the top 20 PCs in each region. PCA was fit separately for N and PN models, and the same PCA loadings learned from training sets were applied to transform test sets. We generated input matrix ***X*** and output matrix ***Y***, where ***X*** can is composed of two submatrices: ***X*** = [***X***_*d*_ ***X***_*u*_]. We predicted ***Y*** using linear regression:

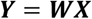

where ***W*** can also be partitioned into submatrices representing upstream and downstream weights as 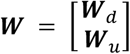. We solved for ***W*** using linear regression with sample weighting (to mediate upweighting) and with Tikhonov regularization:

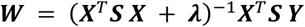

where ***S*** is the sample weighting matrix and ***λ*** is a matrix of regularization strengths. ***S*** is a diagonal matrix of size timesteps x timesteps, whose entries give a sample weight to the activity from each time point. For PN and UN models, we upweighted specific diagonal elements (corresponding to specific timesteps) to 50, corresponding to 50x weight in the model, while all other diagonal elements were set to 1. For PN models, we upweighted timesteps from perturbation segments, and for UN models, we upweighted timesteps from ‘upweight segments’ (see above).

The regularization matrix ***λ*** was partitioned into block submatrices to apply penalties separately to upstream and downstream components.

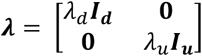

where *λ*_*d*_ and *λ*_*u*_ are scalars that penalize the corresponding downstream and upstream weights, and ***I***_***d***_ and ***I***_***u***_ are identity matrices with size corresponding to the number of regressors from the downstream and upstream regions, respectively.

#### Model evaluation

We evaluated the prediction for each downstream PC using the coefficient of determination:

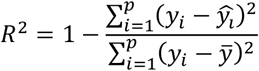

where p is the downstream PC being predicted. As this metric has a range of [-∞, 1], and a single outlier value can skew averages, we defined *R*^2^’s below -1 to be -1. We weighted this *R*^2^ by the variance of the prediction of each PC and then averaged across all 20 PCs, yielding a weighted *R*^2^ value.

#### Region quotient calculation

We calculated region quotients (RQs) by summing the 1-norm of the upstream and downstream contribution for each predicted PC.

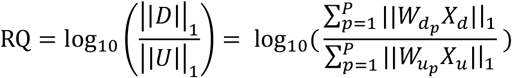

where *p* is the downstream PC being predicted, and 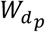 and 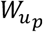 are the learned weights of the downstream and upstream predictions of *p*.

#### Regularization sweeps

To generate models spanning a range of RQs, we repeatedly fit models while varying *λ*_*d*_ and *λ*_*u*_. Intuitively, when *λ*_*d*_ ≫ *λ*_*u*_, it encourages models to have stronger upstream weights *W*_*u*_ than downstream weights *W*_*d*_, and visa versa when *λ*_*d*_ ≪ *λ*_*u*_. We chose pairs of *λ*_*d*_ and *λ*_*u*_ values from the Cartesian product of two vectors of 18 logarithmically spaced numbers between 10 and 10, fitting a model for each pair of values (18^2^ = 324 total models). We discarded all models with an RQ less than -2 or greater than 2.

For the example visualizations of model performance versus RQ (Fig. 2a,b and Fig. 3c,d), we showed only the best-performing model within 12 equally spaced bins of RQ values. For the analysis quantifying the range of well-fit models and the amount of unique information (Fig. 2c,d and Fig. 3f,g), we analyzed all 324 fit models. For the analysis quantifying unique information (Fig. 2d and Fig. 3f), we computed the *R*^2^ of a downstream-only model by fitting a model that only used downstream PCs as regressors. For the analysis of best-fit model predictions (Fig. 2e and Fig. 3h), we fit models using a standard ridge regularization approach where *λ*_*d*_ = *λ*_*u*_. We determined an optimal value for *λ* by sweeping 10 logarithmically spaced values of *λ* between 10 and 10 and evaluating performance on an additional 20% of held-out data.

### Nonlinear Modelling

*Model details:* Here, we predicted the single-neuron downstream firing rates from the past firing rates of downstream and upstream neurons. To do so, we averaged activity in 10 ms bins, and then constructed ***X***_*u*_, ***X***_*d*_, and ***Y*** using single-neuron firing rates.

We predicted activity ***Y*** as:

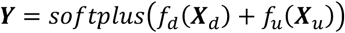

where *f*_*d*_ and *f*_*u*_ are separate learned feedforward neural networks (FNNs) that act on the downstream and upstream activity respectively. The softplus ensures that predicted rates are positive.

We fit the parameters of the downstream and upstream neural networks, which we will refer to as *W*_*d*_ and *W*_*u*_, by minimizing the negative Poisson log-likelihood of the predicted activity, with L_2_ regularization and sample weighting:

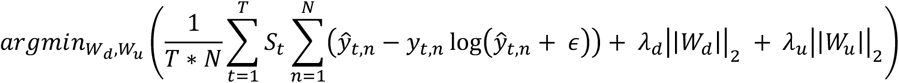

where *N* is the number of downstream neurons, *T* is the number of timesteps, *S*_*t*_ is the sample weight at time *t*, and *λ*_*d*_ and *λ*_*u*_ are regularization strengths for downstream and upstream regions, respectively.

For PN and UN models, we upweighted specific timesteps by setting specific *S*_*t*_ values to 30, corresponding to 30x weight in the model, with all other elements set to 1. We empirically found that sample weights higher than 30 tended to cause overfitting to the upsampled time points. For PN models, we upweighted perturbation segments, and for UN models, we upweighted designated upweight segments (see above). Sample weight values were chosen to maximize predictivity on an additional held-out 20% of data (‘validation set’).

Each FNN consisted of 3 layers with 200 hidden units per layer. We set the learning rate to 10^−3^ and learned weights using the Adam optimizer. Training ran until the loss on the validation set stopped decreasing, or when 1000 iterations was reached.

*Regularization sweeps:* In order to learn a range of models that spanned RQs between -2 and 2, we varied pairs of *λ*_*d*_ and *λ*_*u*_ terms algorithmically. To begin sweeps, *λ*_*d*_ was initialized with a high regularization term (10^0^ for most sessions) and *λ*_*u*_ with a low regularization term (10^−2^ for most sessions), and *λ*_*d*_ was progressively decreased at an adaptive rate to generate models with increasing RQs. Once *λ*_*d*_ reached the low regularization term value, *λ*_*u*_ was progressively increased at an adaptive rate to continue generating models with increasing RQs. Sweeping was stopped once models spanned RQs from -2 to 2 and all adjacent models differed by less than 0.15 RQ.

Models with very similar RQs tended to vary substantially in terms of their predictivity, likely due to the stochastic nature of fitting neural network models (unlike linear models). To account for this stochasticity and ensure a smoothly varying performance across the range of RQs, we divided up the RQs in 30 overlapping bins and took the median predictivity for all models within that bin. This resulted in 30 final models, equally spaced apart, spanning RQs from -2 to 2. For the example visualizations of model performance versus RQ (Fig. 2a,b and Fig. 3c,d), we showed only 12 of these models. For the analysis quantifying the range of well-fit models and the amount of unique information (Fig. 2c,d and Fig. 3f,g), we analyzed all 30 models. For the analysis quantifying unique information (Fig. 2d and Fig. 3f), we computed the *R*^2^ of a downstream-only model by fitting a model that comprised only of a downstream FNN. For the analysis of best-fit model predictions (Fig. 2e and Fig. 3h), we fit models using a standard regularization approach where *λ*_*d*_ = *λ*_*u*_. We determined an optimal value for λ by sweeping 10 logarithmically spaced values of λ between 10^−2^ and 10^0^ and evaluating performance on the validation set.

#### Region quotient calculation

We calculated the region quotient by summing the 1-norm of the output of the downstream and upstream FNN for each predicted neuron:

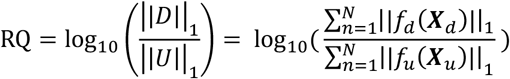

where N is the number of downstream neurons being predicted, and *f*_*d*_ and *f*_*u*_ specify the downstream and upstream FNN functions.

Model *evaluation:* We evaluated the goodness-of-fit of predictions for neurons’ spike counts using pseudo-*R*^2^ (a generalization of *R*^2^ for non-Gaussian distributions)^25^:

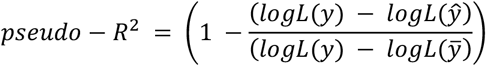

where *logL*(*ŷ*), *logL*(*y*), and 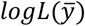 are the Poisson log-likelihoods of predicting the true spike count of the data according to the predicted rate *ŷ*, the true spike counts *y*, and the mean spike count 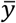, respectively.

### Multi-region RNN Simulations

*Overview:* To generate simulated data where the ground truth interaction between regions was known, and with data comparable to our recordings, we took the approach of first fitting multi-region RNNs to our naturally occurring recorded data. We then simulated data from these RNN models. We chose one session from each of three mice, and then fit two networks per session, yielding six total RNNs.

#### RNN fitting

To preprocess the firing rate time series for fitting with multi-region RNNs, we removed all non-climbing periods and excluded all neurons with an average firing rate under 0.5 spikes per second. We averaged firing rates in 5 ms bins and then smoothed using a Gaussian filter with a standard deviation of 50 ms. After smoothing, we z-scored the firing rate of each neuron and then divided by the population-wide maximum value.

We fit RNNs using a modified version of the CURBd^18^ Python package. Here, a multi-region RNN contains a number of units equal to the number of recorded neurons, and the activity in each unit evolves according to the following equation:

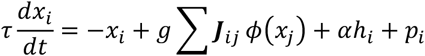

where *τ* is the decay constant in seconds, *dt* is the length of each timestep, *x* is the activity of target unit *i, x*_*j*_ is the activity of source unit *j, g* scales the strength of recurrent connections, ***J*** is a learned weight matrix denoting the recurrent weights from source units *j* to target units *i, ϕ* is a *tanh* activation function, *h* is a white noise external input scaled by α, and *p* is an additional external input we used in our simulations to induce perturbations.

We modified the CURBd training function to ensure across-region weights were sparse and positive. To do so, we first modified the random initialization of the ***J*** weight matrix such that all across-area weights were positive, sampled from *N*(0,1) with negative values omitted. To better ensure training stability with this modification, we also subtracted all within-area weights by a small scalar value during initialization.

Then, every training epoch, we set to 0 any across-area weights that fell below 0. To ensure sparsity, we ranked neurons by the 1-norm of their across-region weights, and then set the bottom *r* percentile of neurons to have across-region weights of 0. We initialized *r* to 50% of its final value at the start of training, and gradually increased *r* over the course of training to its final value (which varied by session; see below). All neurons that ended training with all across-area weights of 0 we considered “interneurons”. We discarded and replaced the few fit models where these interneurons did not have mostly inhibitory weights.

We used standard fitting parameters recommended for electrophysiological data in CURBd. We set *τ* to 0.05, *dt* to 0.001, α to 0.001, the learning rate to 1.0, and the initial value of *g* to between 1.4 and 1.6, differing slightly across the six networks. To span a range of relative across-region connectivity across the six different RNNs, we varied *r*, the sparsity percentage, across the six RNNs from 0.3 to 0.7, such that RNNs with high *r* (less across-region connections) had generally weaker across-region connectivity. During fitting, the 100 timesteps were iteratively computed before each model evaluation. We trained for either 10 or 15 training runs, which was when R^2^ on training data no longer substantially increased with additional runs.

#### Simulations

We used RNNs to run simulations with sporadic activity perturbations driven by adding input to the upstream “interneurons”. We randomly initialized the RNN with an activity state randomly sampled from those in the training data. We let the simulation run for 30,000 timesteps to allow stabilization, then induced perturbations every 2.5 seconds with 25-ms of positive input *p* to all “interneurons” in the upstream region. This input was sampled from a *N*(0,3*α*) with negative values omitted, such that the perturbation input was 3x the variance of *h* . This input created short-latency suppression in the upstream region and then subsequent suppression in the downstream region, matching the approximate time course of perturbations in actual recordings. We discarded RNNs whose perturbation segments did not resemble those in actual recordings. We ran simulations for a total of 300,000 timesteps.

The simulation outputs were normalized rates between -1 and 1 for every neuron. To sample spike trains from these underlying rates, we first needed to convert these normalized rates back into typical firing rate space. To do so, we first rescaled rates by the population-wide maximum value of the original z-scored firing rates. Then, we added the overall mean and scaled by the overall standard deviation of the original firing rates to reverse the z-scoring. We set all negative firing rate values to 0 and multiplied all firing rates by 5, an arbitrary scale factor needed to ensure model performance was comparable to fits of actual data. We then generated spike trains by taking Poisson samples from these underlying firing rates.

#### Simulated data for modelling

For the RNN-simulated data that we fit using our linear and nonlinear models, we extracted perturbation segments and naturally occurring segments. Perturbation segments were defined as the -50 to 40 ms window around perturbation onset. We note that using the -50 to 70 ms window used in actual recordings slightly increased prediction errors here; we used the longer time window in analyses of actual recordings to maximize data for fitting given the limited number of optogenetic perturbations per session. We held out naturally occurring segments (‘control segments’) from randomly sampled 90 ms random windows within the 400-1000 ms after the onset of each perturbation. Naturally occurring segments consisted of all other neural activity beginning at least 400 ms after each perturbation onset. Within naturally occurring segments, we also defined randomly chosen 90 ms windows, equal in number to perturbation segments, as upweight segments.

#### Model details

All linear and nonlinear model analyses matched those performed for actual recordings.

#### True RQ calculation

The interaction matrix ***J*** can be partitioned into submatrices ***J***_*u*_ and ***J***_*d*_, where 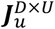 is a weight matrix of size D downstream neurons x U upstream neurons that reflects the strength of upstream input to downstream neurons, and 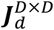 is a square matrix reflecting the strengths of connections between downstream neurons.

The activity of downstream neurons evolves over time according to:

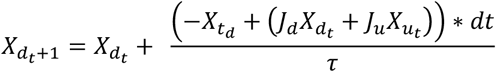

We can arrange this into separate terms:

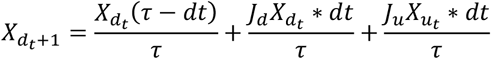

This equation can be rewritten with the terms representing the self-decay, the downstream contribution, and the upstream contribution respectively:

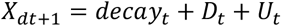

where *D*_*t*_ and *U*_*t*_ are the downstream and upstream contribution at time *t*.

To calculate the true RQ, we sum the 1-norm for each timestep, including the self-decay term as part of the downstream component, as these self-terms are included in the downstream component of the linear and nonlinear models we fit. We set *dt* as 0.01, corresponding to the 10 ms bins used to average activity.

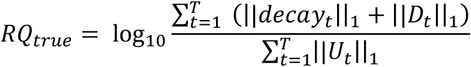

where *T* is the number of timesteps in the simulations.

For PN and N models, we calculate this true RQ based on the simulated activity in the held-out data, which differed between the two model types. Thus, the true RQ from the same network differed slightly between the model types.

## Author Contributions

D.B., A.M., and J.G. devised the project and designed experiments. A.K., D.B., and P.W. performed experiments. D.B. and J.L analyzed the data. D.B., A.M., and J.G wrote the manuscript.

## Acknowledgements

We are grateful to Northwestern’s Quest High-Performance Computing Cluster for enabling access to computational resources. A.M. was supported by a Searle Scholar Award, a Sloan Research Fellowship, a Whitehall Research Grant Award, The Chicago Biomedical Consortium with support from the Searle Funds at The Chicago Community Trust, the Simons Foundation, and NIH grant DP2 NS120847. A.K. was supported by the Dr. John N. Nicholson Fellowship and NIH training grant 2T32MH067564. J.G was supported by NIH grant R00 NS119787 and acknowledges support from the National Institute for Theory and Mathematics in Biology through the National Science Foundation (DMS-2235451) and the Simons Foundation (MPTMPS-00005320).

## Materials & Correspondence

Correspondence and request for materials should be addressed to D.B., A.M., and J.G.

## Data and code availability

All code for data analyses will be made available on Github upon publication. The data that support the findings of this study will be available from a public data archive upon publication.

